# Meiofauna is an important, yet neglected, component of biodiversity of *Posidonia oceanica*

**DOI:** 10.1101/2021.05.09.443247

**Authors:** Guillermo García-Gómez, Álvaro García-Herrero, Nuria Sánchez, Fernando Pardos, Andrés Izquierdo-Muñoz, Diego Fontaneto, Alejandro Martínez

**Affiliations:** Department of Biodiversity, Ecology and Evolution, Department of Biology, Universidad Complutense de Madrid, C/José Antonio Novais 2, 28040, Madrid, Spain; Molecular Ecology Group (MEG), Water Research Institute (IRSA), National Research Council of Italy (CNR), Largo Tonolli 50, 28922, Verbania, Italy; Department of Earth, Oceans and Ecological Sciences, School of Environmental Sciences, University of Liverpool, 4 Brownlow Street, L69 3GP, Liverpool, United Kingdom; Marine Biology Laboratory in Santa Pola (CIMAR), Marine Research Center of Santa Pola, University of Alicante, Aptdo. 99, 03080, Alicante, Spain

**Keywords:** Acari, Biodiversity, Copepoda, Habitat sorting, Nematoda

## Abstract

*Posidonia oceanica* meadows are biodiversity reservoirs and provide many ecosystem services in coastal Mediterranean regions. Marine meiofauna, on the other hand, not only represents a major component of regional marine biodiversity, but also a useful tool to address both theoretical and applied questions in ecology, evolution, and conservation. We review the meiofaunal diversity in the *P. oceanica* ecosystem combining a literature review and a case study. First, we gathered records of 664 species from 69 published studies as well as unpublished sources, including few species exclusive from this ecosystem. Eighteen of those studies quantified the spatial and temporal changes of species composition, highlighting habitat-specific assemblages that fluctuate following the annual changes experienced by *P. oceanica*. Hydrodynamics, habitat complexity, and food availability, all three inherently linked to the seagrass phenology, are recognised as the main factors at shaping the complex distribution patterns of meiofauna in the meadows. These drivers have been identified mainly from Copepoda and Nematoda, and depend ultimately on species-specific preferences. Second, we tested the generality of these observations using marine mites as a model group, showing that the same processes might be in place also for other less abundant meiofaunal groups. Overall, our study highlights an outstanding diversity of meiofauna in *P. oceanica* and shows its potential for future research, not only focused on exploring and describing new species of neglected meiofaunal organisms, but also providing a more complete understanding on the functioning of the iconic Mediterranean ecosystem created by *P. oceanica*.

## 1. INTRODUCTION

Named after the Greek God Poseidon, the marine flowering plant *Posidonia oceanica* (L.) Delile, 1813 constitutes an iconic Mediterranean endemic organism that forms extensive lush meadows, imprinting shallow Mediterranean bays and beaches with a unique character. Beyond the ethnographic importance, meadows of *P. oceanica* also provide many ecosystem services. The leaves, which may extend around one meter beyond the seafloor, act as a major carbon sink filtering and oxygenating the seawater (Bay, 1984; Pirc, 1985; Mateo et al., 1997; Terrados & Duarte, 2000), and shelter the local hydrodynamics favouring sedimentation (e.g., Gacia & Duarte, 2001; Folkard, 2005; Manca et al., 2012). Underneath the leaves, a monumental formation of rhizomes, roots, and detritus, typically termed as “matte” (Boudouresque & Jeudy de Grissac, 1983), stabilises the sediment and entraps particles of organic matter (Mateo et al., 1997; Pergent et al. 2012). Leaves and matte together prevent erosion in the littoral zone, support food webs, and enrich the surrounding bare sand with organic matter and nutrients (Jørgensen et al., 1981; Simeone & De Falco, 2013; González-Ortiz et al., 2014; Connolly & Waltham, 2015). The adjacent sandy areas form corridors and wide spaces among seagrass patches, constituting an ideal zone at the meadow’s edges, where certain species settle (Coppa et al., 2010), seek refuge (Pinna et al., 2013), and feed (Sánchez-Jerez et al., 1999a). Furthermore, exported materials from the meadows accumulate and constitute other habitats, mostly on nearby sandy areas (Dimech et al., 2006; Cresson et al., 2012), but also further down to deep-waters or into caves and pits (Picard, 1965), where they boost local food webs as an additional source of allochthonous nutrients (Romero et al., 1992; Mateo et al., 2003; Guala et al., 2006; Simeone et al., 2013). Even when washed-up on the seashore, *P. oceanica* detritus cover vast extensions with complex dynamics throughout the year, called “banquettes” (Boudouresque & Meinesz, 1982). Banquettes are fundamental in protecting beaches from erosion, stabilising the sand dune, and enriching underlying sediments with nitrogen (Boudouresque et al., 2016). Due to such significant and multifaceted modification of the environment, *P. oceanica* is considered as an ecological engineer (Unsworth & Cullen-Unsworth, 2017), which forms a complex ecosystem composed of living seagrass and its exported detritus, as well as the rich community of organisms associated to its different habitats (Mazzella et al., 1989; Boström et al., 2006).

Over the last decades, scientists have described the diversity and functioning of the *P. oceanica* ecosystem, along with the local dynamics of the mosaic of habitats associated with them. Initial research focused on the phenological annual changes of the *P. oceanica* plants themselves (e.g., Ott, 1980; Bay, 1984), followed by detailed characterizations of the diversity and structure of their associated communities. Indeed, many studies have addressed the dynamics of *P. oceanica* meadows highlighting the economic and ecological importance of their inhabitants (e.g., Dimech et al., 2002; Duffy et al., 2003; Como et al., 2008; Honkoop et al., 2008; Kalogirou et al., 2010; Whippo et al., 2018). For instance, the leaves foster nurseries of fish and cephalopods that often represent important resources for local fisheries (Cetinić et al., 1999, 2011), whereas the assemblages of gastropods and bivalves associated to the plant include several emblematic, often endemic, Mediterranean species (Urra et al., 2013). In the matte, many species of infaunal crustaceans, molluscs, and annelids thrive (Borg et al., 2006), contributing to the overall recycling of the entrapped organic matter (Vizzini et al., 2005). Even beyond the plant, the phytal detritus drifting away from the meadows on the adjacent sandy areas host diverse macrofaunal communities (Sánchez-Jerez et al., 1999b; Guidetti, 2000; Gallmetzer et al., 2005). However, the epiphytic organisms growing throughout the plant structure constitute the communities that have received most attention, partially due to their high species diversity (ca. 660 species, after Piazzi et al., 2016), but also because they have been used as bioindicators of the health of the meadows (Martínez-Crego et al., 2010; Giovannetti et al., 2010; Mateu-Vicens et al., 2014). Interestingly, such research on epiphytic organisms has shown that the distribution of these species is not uniform within the seagrass, but rather the opposite: some epiphytes dominate the older but more exposed leaf tips, others prefer their sheltered middle or basal parts, and few even select the shaded rhizomes (Gambi et al., 1995; Piazzi et al., 2016).

In contrast with macrofaunal communities, information is much more scattered for the meiofaunal animals inhabiting the P. oceanica ecosystem. Meiofauna includes a heterogeneous subset of organisms that are retained between a tandem of meshes with 500 µm and 63 µm size, largely dominated by microscopic animals, but also including larger forms with elongated morphologies or contractible bodies (Giere, 2009; Schmidt-Rhaesa, 2020). Meiofauna plays a fundamental role in many processes in the seafloor (Schratzberger & Ingels, 2018) and represent a numerically important, yet often neglected, component of the diversity of many regions (Curini-Galletti et al., 2012, 2020; Martínez et al., 2019). Furthermore, meiofauna represents an invaluable tool to address both theoretical and applied eco-evolutionary questions (e.g., Fontaneto et al. 2007; Fontaneto 2011; Gansfort,; Laumer et al. 2015; Martínez et al., 2020; Fontaneto & Zhai, 2020; Martin-Duran et al., 2021), as these organisms belong to virtually every animal phylum, thus alleviating the confounding effect of potential phylogenetic bias during inductive hypothesis tests (Giere, 2009; Rundell & Leander, 2011). Moreover, meiofauna critically supports marine food webs by transferring the energy from decomposer and primary producer microorganisms to higher trophic levels (Danovaro, 1996; Danovaro et al. 2007). Although scattered in the literature, numerous records indicate that many meiofaunal species inhabit the *P. oceanica* ecosystem, whether crawling on the leaves or across the matte labyrinths, gliding in the interstices of the adjacent sediments, or even drifting within the detritus over the sea bottom. Several studies addressed the composition and community dynamics of some meiofaunal groups in *P. oceanica*, revealing that the biotic and abiotic conditions of the meadow shape the distribution of meiofaunal species (e.g., Novak, 1982, 1989; Mascart et al., 2013; Mascart, Lepoint, et al., 2015; Pusceddu et al., 2016). However, besides these few comprehensive ecological studies, mainly focusing on copepods and nematodes, most of this research involves occasional taxonomic descriptions from punctual samples. Unfortunately, the lack of a comprehensive updated review of all this literature obscures our understanding of the overall diversity and role of meiofauna in this iconic Mediterranean ecosystem.

The aim of this study is two-folded. First, we reviewed the diversity (i.e., species richness) and ecology of meiofauna in *P. oceanica* through a literature survey, completed with unpublished data provided by our colleagues. Despite our aim is mainly exploratory (Yanai & Lercher, 2020), we depart from the assumption that the patterns in meiofaunal diversity in *P. oceanica* resembles those described for larger organisms. We expect that meiofaunal species do not occur homogeneously in the *P. oceanica* ecosystem, but they segregate spatially and temporally across its different habitats following the annual phenological cycle of the meadows. Second, we selected a case study to test explicitly the latter assumption. We used halacarid marine mites as model organisms because they are common in seagrasses such as *P. oceanica* (Mari & Morselli, 1990; Zupo, 1993; Durucan, 2018; Durucan & Boyacı, 2018; Durucan & Levent, 2019), yet the habitat preferences of the species associated with this plant have never been explicitly investigated. This case study, therefore, provides a point of comparison with the better studied copepods and nematodes (Novak, 1989; Mascart et al., 2013)

## 2. MATERIALS AND METHODS

### 2.1 Literature review

#### 2.1.1 Selection of references

We systematically screened Google Scholar for all published literature containing records of vagile meiofaunal species within *Posidonia oceanica* habitats. The search, performed in December 2020, consisted of the word ‘Meiofauna’ or the name of a target animal group (i.e., ‘Acari’, ‘Annelida’, ‘Cephalocarida’, ‘Copepoda’, ‘Gastrotricha’, ‘Gnathostomulida’, ‘Kinorhyncha’, ‘Loricifera’, ‘Mystacocarida’, ‘Nematoda’, ‘Platyhelminthes’, ‘Rotifera’, ‘Tardigrada’) followed by the term ‘Posidonia’ (e.g., ‘Copepoda’ AND ‘Posidonia’). After screening the abstract of all the compiled references, all papers including relevant information were downloaded. To maximize the completeness of our database, the references cited in all downloaded papers were checked for additional sources.

#### 2.1.2 Compilation of the species inventory

First, to evaluate the known diversity of meiofaunal species in the *P. oceanica* ecosystem, we carefully screened each selected paper to compile all available records. Each entry included the taxon name, locality, WGS84 geographic coordinates (extracted directly or calculated after the description of each locality), depth, collection method, and the habitat within the *Posidonia oceanica* ecosystem, when these data were available (see Supplementary Information Table S1). We considered four types of habitats: (1) seagrass, consisting of the structure created by the plant; (2) adjacent sediments, including the bare sediments and interstitial habitats next to seagrass patches; (3) macrophyte detritus, the wandering vegetal debris from *P. oceanica* seagrass accumulated on sediments (termed as ‘macrophytodetritus accumulations’ in Mascart, Lepoint, et al., 2015); and (4) banquette, the detritus deposited on the shore after the falling of the *P. oceanica* leaves (Boudouresque & Meinesz, 1982). When this information was provided, we further divided the seagrass habitat in its two discrete compartments: the leaves and the matte, i.e., the underlying ensemble of living rhizomes, plant debris, and roots (see Figure 2). This information was subsequently used to evaluate the habitat preferences of the meiofaunal species living in the *P. oceanica* ecosystem (see below). Furthermore, to maximize the number of provided records, we consulted different specialists, who kindly shared their unpublished records to this study.

#### 2.1.3 Review of ecological questions

Complementary to the inventory of species, all papers with an ecological scope in our database were further screened for relevant hypotheses and findings regarding the preferences of meiofaunal species within the habitats of *P. oceanica*. The information contained in these papers was organised in a table (Table S2), comprising the aim of the study, the targeted meiofaunal groups, and the diversity metrics implemented. For each study we further provided a concise summary of the findings, on whether they found differences in the biodiversity metrics or abundance between (i) habitats, (ii) between samples within the same habitat from a given locality (i.e., local scale), (iii) between localities (i.e., regional scale), and (iv) over time. Last, the table included the future research questions proposed by each study.

### 2.2 Case study: halacarids in *P. oceanica*

#### 2.2.1 Study site and sampling design

In addition to our literature survey, we investigated the habitat specificity in the community of halacarids inhabiting a *Posidonia oceanica* meadow in Cala del Cuartel (Alicante, SE Spain; WGS84 coordinates 38° 12’ 34.04’’ N, 0° 30’ 19.12’’ W), located in a region where animal communities associated to these meadows have been well documented (Villora-Moreno et al., 1991, 1997; Sánchez-Jerez et al., 1999a, 1999b; Martínez, García-Gómez, in press). Sampling was carried out during four campaigns in December 2015, and March, April and August 2016, each coinciding with a different season. During each campaign, SCUBA divers sampled six randomly selected patches of *P. oceanica*, totalling 24 patches for the whole study. Samples of (1) leaves, (2) matte, and (3) adjacent sediments were collected at each patch and standardised through 400 cm^2^-quadrats (20 × 20 cm) (e.g., Novak, 1989; Sánchez-Jerez et al., 1999a; Pusceddu et al., 2016; but see Bell et al., 1984). The leaves were first cut at the ligula level and collected carefully with a hermetic bag; then, the underlying matte was shovelled into another hermetic bag (following Novak, 1982, 1989; Cvitković et al., 2017). Identical samples of the adjacent sediments to each sampled seagrass patch were collected using jars.

Halacarid mites were extracted combining magnesium chloride and “bubble and blot” techniques (Higgins & Thiel 1988, Sørensen & Pardos 2008), filtered through a 63 μm mesh, and fixed in 7% formaldehyde. Fixed halacarids were then sorted using a MOTIC® SMZ-168 stereoscope and whole-mounted on slides in a modified Hoyer’s medium (Mitchell & Cook 1952). Whole-mounted specimens were examined using an Olympus DP70 camera mounted on a light microscope equipped with differential interference contrast microscopy (DIC). We followed André (1946), Green & MacQuitty (1987), and specific taxonomic literature (Morselli, 1980; Bartsch, 1986, 2001, 2006) for species identification. Adult and juvenile specimens were distinguished following Bartsch (2015). Nomenclature followed the World Register of Marine Species (WoRMS Editorial Board 2021).

#### 2.2.2 Data analysis

All statistical analyses were performed using the *R* software version 3.6.1 (R Core Team 2019). We investigated the variation in species richness (i.e., number of species), abundance (i.e., number of individuals) and evenness (i.e., Pielou’s J) within the leaves and the matte, because no mites were found in the sediments adjacent to the seagrass. First, we investigated differences in species richness, abundance (as log_10_ to cope with stark differences between samples), and evenness between leaves and matte samples collected from the same sampling point, using a paired-samples t-test (paired t-test) through the function ‘t.test’. Pielou’s J was calculated for each sample using function ‘diversity’ in the R package *vegan* v. 2.2-1 (Oksanen et al., 2015) to obtain first the Shannon index and then dividing it by the natural logarithm of the number of species. For one sample in the matte, Pielou’s J was not calculated as only one species was observed. Second, we tested whether richness or abundance changed with food availability and habitat complexity, as well as over time. We used the length of the leaves and the organic carbon content of the sediment as a proxy for food availability in the leaves and the matte, respectively. The length of the leaves was measured as the average distance in centimetres from the ligula to the apical end of all the complete leaves found in each sample, which is known to correlate positively with the abundance of epiphytic microorganisms (Mabrouk et al., 2010) that may serve as food for many halacarid species (Bartsch, 1989). Likewise, the percentage of organic carbon of the sediment was calculated using the Walkley-Black method (Walkley & Black, 1934), indicating the amount of accumulated organic matter in the matte, representing available food for mites. Habitat complexity was inferred through the density of leaves or matte, calculated as the dry weight of the leaves or the matte divided by the total volume of the habitat, which varied in the leaves (Average leave length * 20 cm x 20 cm) and was constant in the matte (2 cm x 20 cm x 20 cm). We performed linear models using function ‘lm’ to examine the effect on richness, abundance (as log_10_), and evenness of the environmental variables within each habitat: the length and density of the leaves within the leaves, and the percentage of organic matter and density of the matte within the matte. We accounted for the temporal variation in the seagrass over the period of study by including the sampling date in all models. The significance of each independent variable was summarized as a Type II ANOVA table, using the function ‘Anova’ in the R package *car* version 3.0.9 (Fox et al., 2012). The assumptions of the linear models were controlled visually by checking the normality of model residuals, the plots of residual versus fitted values, and normal Q-Q plots (Crawley, 2013).

To further investigate whether halacarids preferred a certain habitat, we performed two additional analyses. First, since different ecological preferences have been reported between life stages in halacarids (Somerfield & Jeal, 1995, 1996; Bartsch, 2004), we tested for differences in percentage of juveniles (i.e., number of non-adult individuals / total individuals of each sample; in %) between the matte and the leaves within sampling points. Second, we tested for differences in abundance between habitats of the dominant species found within each habitat, for which data were sufficient to perform the analyses. Again, we performed paired t-tests for both differences in percentage of juveniles and abundance of the dominant species between habitat samples within sampling points. Paired samples were removed from the analyses when no individuals were found in a sampling point. For all paired t-tests, we first checked the normality of the differences between paired samples by Shapiro-Wilk tests using the function ‘shapiro.test’.

Last, we examined the differences in species composition between communities occurring in different habitats including their nestedness and turnover components, through the Jaccard abundance-based index (Legendre, 2014), using the function ‘beta’ in the R package *BAT* v. 1.5.5 (Cardoso et al., 2015). For both species-and community-level tests, the abundances were again transformed to log_10_ (abundance + 1) to cope with stark differences in abundance between samples as well as with 0 values. We then assessed the percentage of variability in community composition observed across samples through a permutational analysis of variance (PERMANOVA) using distance matrices with the function ‘adonis’ included in the R package *vegan*. Again, we explicitly included the sampling date as an independent variable, to account for the temporal variation in the seagrass in structuring differences of species assemblage.

## 3. RESULTS

### 3.1 Review of meiofauna in *P. oceanica*

We reviewed a total of 69 relevant studies: 51 of them consisting of taxonomic papers or species inventories and 18 comprising ecological studies (see Supplementary Information). Together with the data published within the present study, we compiled 1045 records for 664 species (Table S1). Nematoda, Copepoda, and Acari accumulated 73% of the records; Tardigrada, Platyhelminthes, Gastrotricha, and Annelida 21%; Kinorhyncha, Xenacoelomorpha, Ostracoda, Gastropoda, Chaetognatha, Mystacocarida, and Rotifera accounted for the remaining 6% of the dataset. No records were found for Gnathostomulida and Loricifera. The majority of sampling sites were concentrated in the Central Mediterranean Sea, particularly in Corsica, Sardinia, and the Gulf of Naples (Figure 1). Most of the species have been recorded within the seagrass (346 species), mostly in the matte (292 species), as well as in the sediments adjacent to the meadows (275 species) (Figure 2A). Interestingly, ca. 85% of the species recorded in the adjacent sediments and within the seagrass have been exclusively found in these habitats. Copepoda, Platyhelminthes, Gastrotricha, Annelida, Tardigrada, and Xenacoelomorpha were mainly found in the sediments, whereas Nematoda and Acari were observed principally within the seagrass (Figure 2D), being records for Nematoda particularly abundant in the matte. Most studies relied on samples collected by hand (562 species), generally by SCUBA divers (Figure 2B), which recovered generally different species from the studies that relied on boat-operated collections (Figure 2E). Most species and meiofaunal groups were reported from shallow waters (476 species; between 0–20 m), and these numbers decreased with increasing depth (Figure 2C and 2F).

**Figure 1.**
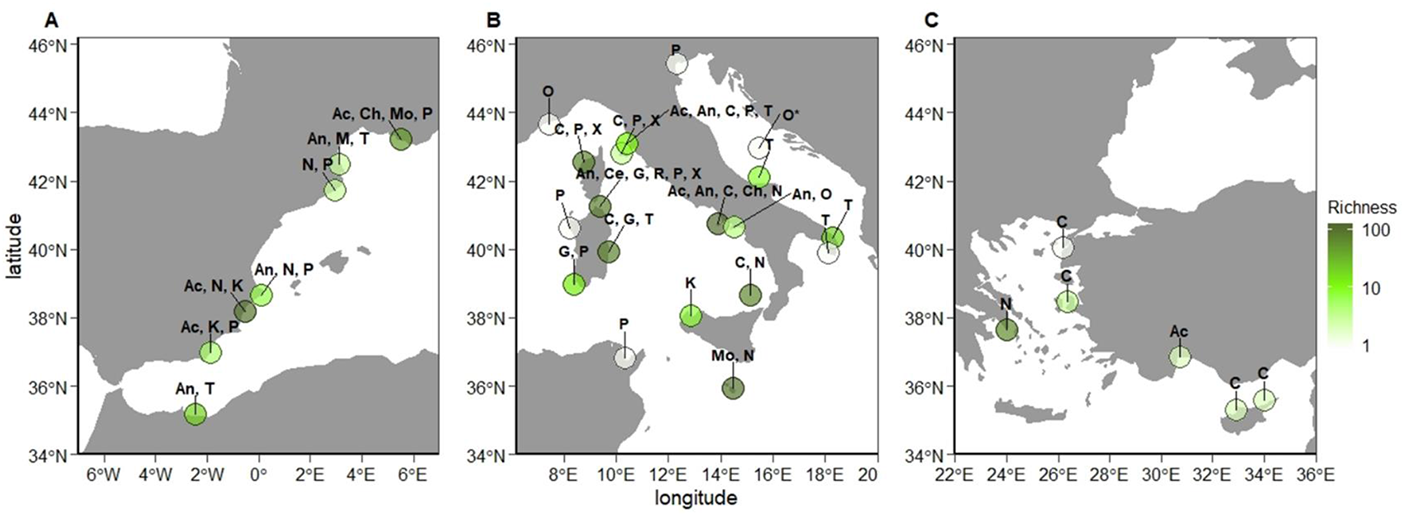
Distribution of the meiofauna recorded in *Posidonia oceanica* in the West (A), Central (B), and East (C) Mediterranean Sea, coloured according to the number of species (see Table S1 for coordinates). Abbreviations: Ac, Acari; An, Annelida, C, Copepoda; Ce, Cephalocarida; Ch, Chaetognatha; G, Gastrotricha; K, Kinorhyncha; M, Mystacocarida; Mo, Mollusca; N, Nematoda; O, Ostracoda; P, Platyhelminthes; R, Rotifera; T, Tardigrada; X, Xenacoelomorpha. *Unspecified location in the Adriatic Sea.

**Figure 2.**
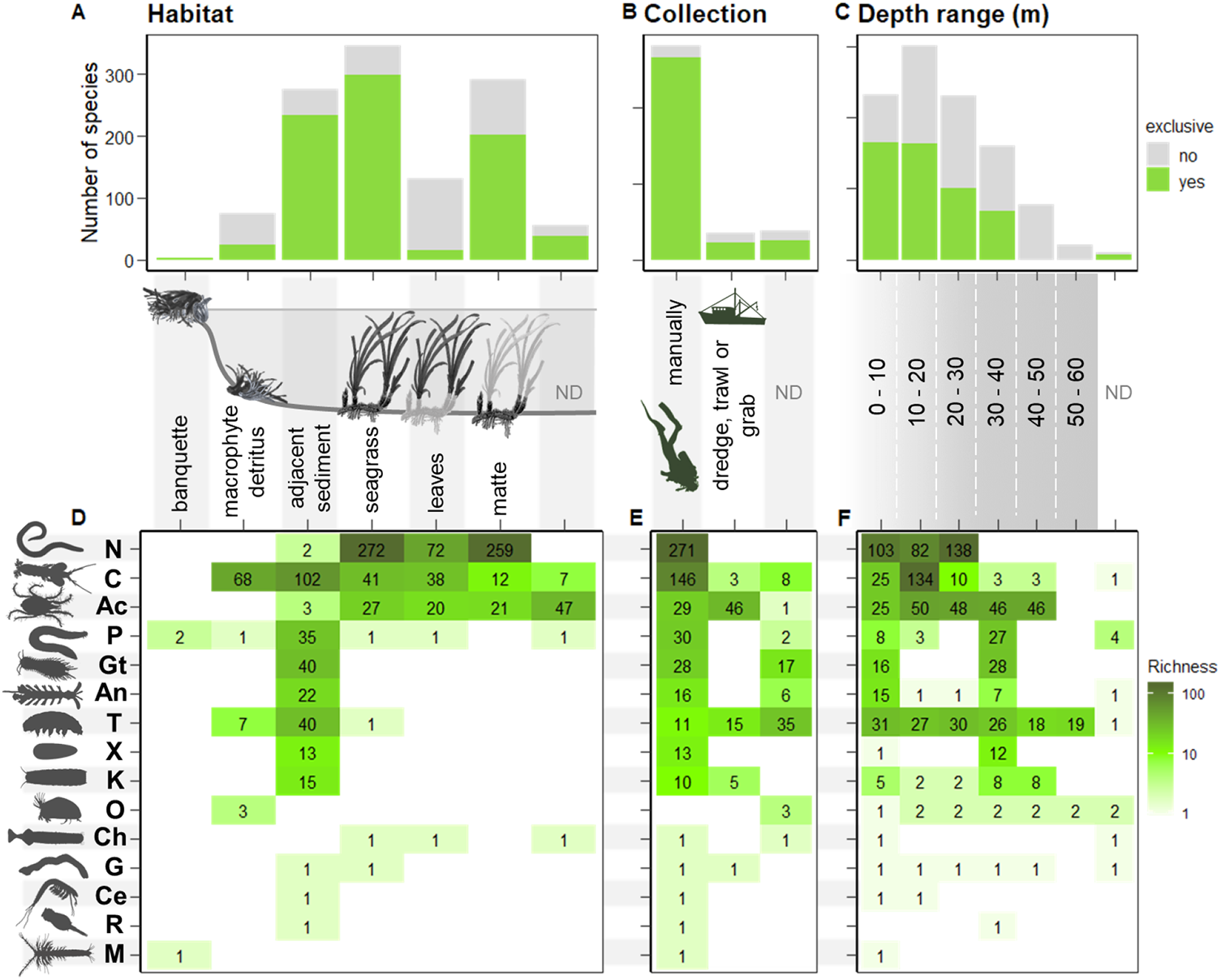
Number of species per meiofaunal group categorized by habitat, collection method, and depth range. Top panels (A-C) indicate the species exclusively observed in one category in green, and those found in more than one category in grey. Bottom panels (D-F) show the number of species per meiofaunal group from light to dark green. In A and D, the seagrass habitat was further divided into the leaves and the matte. Abbreviations as in Figure 1; ND, no data.

We sorted 18 ecological studies (Table S2) that addressed the variation in taxonomic richness, species composition, and abundance of meiofaunal communities at regional (i.e., between localities, generally kilometres away; in nine studies) and local spatial scales (i.e., between samples within the same habitat in a given locality; in 13 studies). Only three of these studies incorporated functional metrics (Novak, 1989; Mirto et al., 2014; Guilini et al., 2017), and a single paper investigated diet preferences (Mascart et al., 2018). Nematodes and copepods were the preferred groups (six studies at specific level), whereas other groups were often recorded at the taxonomic rank of phylum or class. Moreover, 16 studies searched for the ecological factors influencing the presence of meiofauna in different habitats of the meadows, stressing the importance of hydrodynamics, habitat complexity, and food availability. Last, 10 of the studies highlighted temporal variation as another major driver of meiofaunal communities, mainly affecting abundances.

### 3.2 Case study on halacarid mites

We counted 1730 individuals belonging to 21 species and 9 genera (Table 1). One species was restricted to the leaves and four to the matte, whereas 16 species co-occurred in both habitats. No halacarids were found in the sediments. The genus *Copidognathus* was represented by 7 species, followed by the genera *Agauopsis* and *Agaue* (3 species each), and *Rhombognathus* and *Arhodeoporus* (2 species each). The genera *Halacarus*, *Lohmanella*, *Pelacarus,* and *Simognathus* were represented by one species.

**Table 1.** Halacarid species found in this study and their abundances (i.e., number of individuals) and occurrences (i.e., % of samples in which each species was observed) within the leaves and the matte of the *Posidonia oceanica* meadow.

| Species | Abundance |  | Occurrence |  |
| --- | --- | --- | --- | --- |
|  | Leaves | Matte | Leaves | Matte |
| *Agaue adriatica* Viets, 1940 | 2 | 1 | 4.17 | 4.17 |
| *Agaue cf. abyssorum* (Trouessart, 1896) | 0 | 1 | 0.00 | 4.17 |
| *Agaue panopae* (Lohmann, 1893) | 15 | 5 | 41.67 | 12.50 |
| *Agauopsis brevipalpus* (Trouessart, 1889) | 3 | 8 | 12.50 | 29.17 |
| *Agauopsis microrhyncha* (Trouessart, 1889) | 12 | 8 | 45.83 | 25.00 |
| *Agauopsis minor* (Trouessart, 1894) | 13 | 8 | 33.33 | 29.17 |
| *Arhodeoporus gracilipes* (Trouessart, 1889) | 6 | 17 | 25.00 | 54.17 |
| *Arhodeoporus labronicus* (Morselli, 1981) | 0 | 4 | 0.00 | 12.50 |
| *Copidognathus lamelloides* Bartsch, 2000 | 22 | 52 | 41.67 | 66.67 |
| *Copidognathus latisetus* Viets, 1940 | 6 | 20 | 12.50 | 45.83 |
| *Copidognathus magnipalpus* (Police, 1909) | 360 | 23 | 66.67 | 45.83 |
| *Copidognathus oculatus* (Hodge, 1863) | 49 | 9 | 79.17 | 20.83 |
| *Copidognathus quadricostatus* (Trouessart, 1894) | 0 | 3 | 0.00 | 8.33 |
| *Copidognathus remipes* (Trouessart, 1894) | 7 | 21 | 12.50 | 54.17 |
| *Copidognathus reticulatus* (Trouessart, 1893) | 6 | 0 | 8.33 | 0.00 |
| *Halacarus actenos* Trouessart, 1889 | 0 | 1 | 0.00 | 4.17 |
| *Lohmannella falcata* (Hodge, 1863) | 7 | 6 | 20.83 | 16.67 |
| *Pelacarus aculeatus* (Trouessart, 1896) | 1 | 3 | 4.17 | 12.50 |
| *Rhombognathus cf. procerus* Bartsch, 1975 | 0 | 1 | 0.00 | 4.17 |
| *Rhombognathus praegracilis* Viets, 1939 | 901 | 109 | 100.00 | 91.67 |
| *Simognathus minutus* (Hodge, 1863) | 4 | 16 | 12.50 | 37.50 |

Species richness showed no significant differences between habitats (Figure 3A; paired t-test: t = -0.78, *p* = 0.44), though abundances were significantly higher in the leaves than in the matte (Figure 3B; paired t-test: t = 5.77, *p* < 0.001), with 82% of the individuals found in the leaves. Species evenness, however, was significantly higher in the matte than in the leaves (Figure 3C; paired t-test: t = 7.63, *p* < 0.001). *Rhombognathus praegracilis* Viets, 1939 was the dominant (58% of total mites found) and most frequent (96% of total samples) species in the meadow, occurring predominantly in the leaves; after *R. praegracilis*, *Copidognathus lamelloides* Bartsch, 2000 dominated the matte, yet being much sparser in the meadow (4% of total mites). These two species together with *Copidognathus magnipalpus* (Police, 1909) accounted for 85% of the total mite abundance and occurred in more than 50% of the samples. In contrast, the following 8 species ranked by abundance represented only 11% of the total mite abundance, with the remaining species representing only 1%. Three of these rare species were represented by one individual.

**Figure 3.**
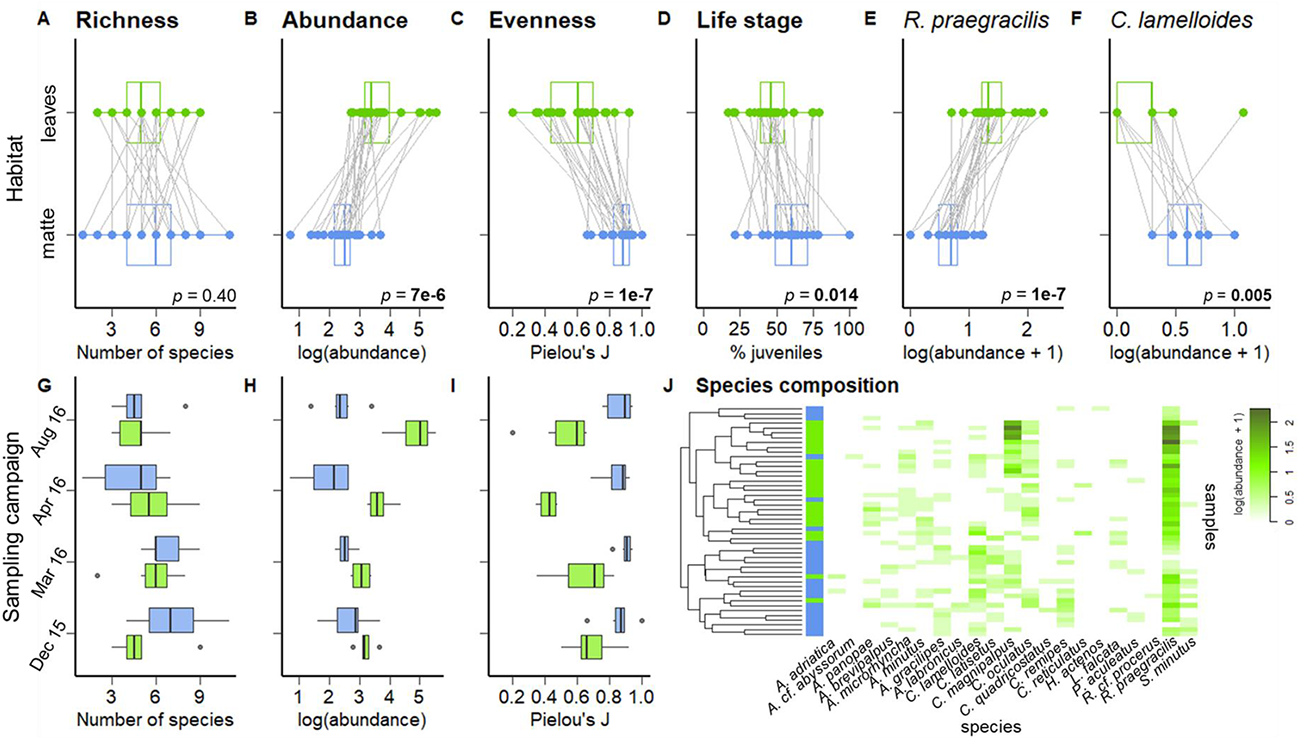
Main findings of the case study on halacarids in a *P. oceanica* meadow (see text for details). Colours differentiate between samples from the leaves (green) and the matte (blue). Top panels (A-F) show paired-t tests and *p*-values for the metrics in this study comparing the leaves and the matte. Panels G-I are boxplots of the variation in species richness, abundance, and evenness in the leaves and the matte through the four sampling campaigns of this study. Panel J show the abundance (as log_10_+1) of each species per sample, coloured from light to dark green. In A-F, samples collected from the same sampling point are connected by lines. In G-I, the error bars indicate the 10^th^ and 90^th^ percentiles of the data, the box’s boundaries indicate the 25^th^ and 75^th^ percentiles, and the solid line within each box marks the median. In J, rows indicate the samples of leaves or matte, whereas columns denote the species; the dendrogram was built using method ‘complete’ by function ‘hclust’. Abbreviations: Dec 15, December 2015; Mar 16, March 2016; Apr 16, April 2016; Aug 16; August 2016.

Halacarid abundances changed significantly over time only in the leaves, reaching the highest abundance in summer, but no temporal change was found in species richness or evenness (Figure 3G-I; Table S3). However, we found no effect of length or density of the leaves on species richness, abundance, or evenness (Table S3). Within the matte, by contrast, halacarid abundance was positively affected by the density of matte (Table S3). No temporal differences (Figure 3G-I) and no effect of any of the environmental variables in this habitat was found on species richness or evenness (Table S3).

Juvenile mites were relatively more abundant in the matte than in the leaves (Figure 3D; paired t-test: t = 2.67, *p* = 0.01). Likewise, we found significant differences in abundance between habitats for the dominant species of the leaves, *Rhombognathus praegracilis*, which preferred the leaves (Figure 3E; paired t-test: t = -7.53, *p* < 0.001), as well as for the second dominant species of the matte, *Copidognathus lamelloides*, which preferred the matte (Figure 3F; paired t-test: t = 3.29, *p* = 0.005). Last, we found that species composition accounting for abundance of mites varied between leaves and matte, through different sampling events in time, as well as with the interaction term between habitat and time (Table 2 and Figure 3J). This suggests that the composition of halacarid species not only differed between habitats, but also shifted within each habitat over the studied period. In addition, most of the differences in species composition were caused by species replacement between habitats (69%).

**Table 2.** PERMANOVA results based on Jaccard dissimilarities using the abundance (as log_10_+1) for the differences in composition of halacarid species between habitats (leaves vs. matte) and through sampling campaigns. *P* values are based on 999 permutations. Abbreviation: Df, Degrees of freedom.

| **Factors** | **Df** | **pseudo-F** | **R<sup>2</sup>** | ***p*** |
| --- | --- | --- | --- | --- |
| Habitat | 1 | 9.958 | 0.164 | **0.001** |
| Sampling campaign | 3 | 1.755 | 0.087 | **0.008** |
| Habitat:Sampling campaign | 3 | 1.793 | 0.089 | **0.004** |
| Residuals | 40 |  | 0.660 |  |

## 4. DISCUSSION

The main findings of the present paper are straightforward: (1) the diversity of meiofauna in *P. oceanica* is not only comparable, but even higher than that reported for other groups of organisms associated with this plant (i.e., epiphytes and macrofauna); (2) most species show specific habitat preferences, which vary from regional to local scales, depending mainly on hydrodynamics, habitat complexity, and food availability. The latter has been shown mainly by copepods and nematodes in the literature, and by marine mites in the presented case study.

### 4.1 Meiofaunal diversity in *P. oceanica*

Overall, we gathered evidence of 664 meiofaunal species occurring in the different habitats provided by the *P. oceanica* (Table S1). Interestingly, 258 of these 664 species were reported by the authors as doubtful or unclear identification (e.g., reporting either ‘sp.’ or ‘cf.’ in the specific name). Since most of these reports were done by internationally acknowledged taxonomical specialists (e.g., Novak, 1989; Curini-Galletti et al., 2012; Guilini et al., 2017), this uncertainty suggests that they may correspond to undescribed, new species to science, or belong to poorly investigated groups meriting further attention. This scenario increases the value of the *P. oceanica* ecosystem as a biodiversity reservoir, not only for macrofaunal species as previously indicated, but also for meiofaunal groups. In fact, the outstanding diversity of meiofaunal species found here largely overcomes the diversity recorded for other groups in species richness. In a similar review, Piazzi et al. (2016) compiled records for 307 species of algae and 353 species of sessile macrofauna in *P. oceanica*, whereas field-based research reports generally lower diversity for other groups. For instance, 68 species of diatoms were reported in a meadow from the Adriatic Sea (Kanjer et al., 2019), 171 species of molluscs in the Alborán Sea (Urra et al., 2013), and 42 species of fish in Hellenic waters (Kalogirou et al., 2010).

Interestingly, meiofaunal species were not reported evenly across the different habitats within the *P. oceanica* ecosystem. While most of the reports correspond to the seagrass habitat (i.e., leaves and matte; 525 records), and the adjacent sediments (341 records),the meiofaunal communities in the macrophyte accumulations (114 records; see also Mascart et al., 2013; Mascart, Lepoint, et al., 2015), and the banquette (6 records; Jansson, 1966; Casu & Curini-Galletti, 2006) remain neglected, despite the latter provides a vast inshore habitat that extends from the water line up to several meters inland (Mateo et al., 2003; Boudouresque et al., 2017). This bias suggests that the actual meiofaunal diversity associated with *P. oceanica* might be much greater than currently reported. This is not a unique feature of these meadows, since meiofauna is often ignored in most biodiversity reports, despite it has been acknowledged as one the major components of diversity in marine ecosystems (Schatzberger et al. 2018). Moreover, many of the species recorded in the literature are putatively exclusive from the *P. oceanica* ecosystem, including evolutionarily interesting species such as the cephalocarid *Lightiella magdalenina* Carcupino et al. 2006, and annelids such as *Psammodrilus curinigallettii* Worsaae, Kvindebjerg & Martínez, 2015 and *P. didomenicoi* Worsaae & Martínez 2018, as well as a single record of *Lobatocerebrum*, possibly corresponding to a new undescribed species (Sanna et al., 2014; Kerbl et al., 2015; Worsaae et al., 2015; 2018).

A similar meiofaunal diversity might occur in meadows dominated by different seagrass species (Sánchez-Jerez et al., 1999a; Cvitković et al., 2017). For example, surveys conducted in meadows of *Cymodocea nodosa* (Ucria) Ascherson, 1870 in the Canary Islands have revealed a large diversity of Gnathostomulida (Sterrer, 1997; Riera, 2012), Chaetognatha (Hernández et al., 2009), Kinorhyncha (Martínez, pers. obs.), and Annelida (Brito et al., 2001, Brito et al., 2005), including the description of new species only known from *Cymodocea* so far (Brito & Nuñez, 2003). These results collectively warrant future comprehensive investigations in other geographical areas and seagrass species.

### 4.2 Habitat preferences in *P. oceanica*

Meiofauna showed differences in habitat occupation in almost all analysed ecological studies (>90%), regardless of the taxonomic ranks of their operational units. Overall, these studies highlighted hydrodynamics, habitat complexity, and food availability as the major drivers for these differences. These drivers not only act hierarchically nested at different spatial scales, but also vary through time. Indeed, the same drivers seem to affect the structure of meiofaunal assemblages in other seagrass ecosystems (mainly copepods and nematodes; see Bell et al., 1984; Decho et al., 1985; Hicks, 1989; De Troch et al., 2001, 2003, 2005, 2006).

At a regional scale, shore hydrodynamics exerts a relatively homogeneous physical pressure on the entire *P. oceanica* ecosystem (Vacchi et al., 2017). However, the great structural heterogeneity of *P. oceanica—*consisting of a mosaic of living plants, macrophyte accumulations, and sandy patches— shelters certain areas within a meadow from the currents (see Abadie et al., 2018). More specifically, patches of high habitat complexity protect the meiofaunal communities from the local hydrodynamics in *P. oceanica* (Mascart, Lepoint, et al., 2015), as commonly observed in seagrasses for other animal groups (e.g., Heck & Orth., 1980; Stoner & Lewis, 1985; Hall & Bell, 1988; Moore & Hovel, 2010). Such sheltering effect is stronger in the matte, which also shows a higher diversity than that in the leaves, both in meiofaunal (Novak, 1982, 1989; Guilini et al., 2017; see Figure 2A and 2D) and macrofaunal species (Gambi et al., 1995; Borg et al. 2006; Piazzi et al. 2016). Food availability within each habitat, finally, drives the presence of different species not only depending on the amount of food (Mirto et al., 2010, 2014; Castejón Silvo, 2011; Losi et al., 2012; Mascart et al., 2013; Cvitković et al., 2017; Polese et al., 2018) but also upon the presence of specific food sources. In fact, it has been shown that even closely related meiofaunal species may prefer different food sources (Mascart et al., 2018).

Temporally, hydrodynamics, habitat complexity, and food availability are inherently linked to the annual cycle of *P. oceanica*, in which long old leaves fall at the end of the summer, being replaced by short young leaves (Larkum et al., 2006). These drastic annual changes in the plant structure affect how the hydrodynamic forces impact *P. oceanica*, since the habitats become increasingly complex and sheltered as the meadow develops (Folkard, 2005). At the same time, densely aggregated leaves foster many epiphytes, offering new and more abundant food sources to the meiofauna (Velimirov & Walenta-Simon, 1993; Mascart et al., 2018; but see Lebreton et al., 2012 on *Zostera noltii*). Indeed, all studies addressing temporal variation in *P. oceanica* found substantial differences over time, not only in taxa composition and abundance (Novak, 1989; Villora-Moreno et al., 1991; Losi et al., 2012; Mascart, Lepoint, et al., 2015; Cvitković et al., 2017; Polese et al., 2018), but also in species’ dietary preferences (Mascart et al., 2018). These studies collectively showed that abundances of most meiofaunal taxa peaked between spring and summer (Novak, 1982, 1989; Villora-Moreno et al. 1991; Sánchez-Jerez et al. 1999a; Losi et al., 2012; Mascart, Lepoint, et al., 2015; Cvitković et al., 2017; Polese et al., 2018), in accordance with the apogee of *P. oceanica*. Similar trends are found in other groups, such as epiphytic forams, diatoms, and dinoflagellates (Piazzi et al., 2016), as well as macrofauna (e.g., Gambi et al., 1992, 1995; Bedini et al., 2011; Urra et al., 2013).

Migration between adjacent habitats is also important to understand the distribution of meiofauna in the meadow (Villora-Moreno et al., 1991; Sánchez-Jerez et al., 1999b; Mascart et al., 2013; Mascart, Agusto, et al., 2015). Meiofaunal migration occurs amongst the different habitats in a meadow and depends on the dispersal ability of the different taxa (Mascart, Agusto, et al., 2015; but see Commito & Tita, 2002; De Troch et al., 2005). These movements may take place seasonally (e.g., nematodes migrating from the matte to the leaves in summer; Novak, 1989), or in shorter time frames (e.g., day-night cycles of vertical migration in copepods; Sánchez-Jerez et al., 1999a). In similar seagrasses, some meiobenthic copepods migrate between the sediment and the vegetated canopy over their life spans (Walters, 1988; Bell & Hicks, 1991), and even within the same day (Hicks, 1986), emerging into the water column at night-time (Bell et al., 1988).

### 4.3 Halacarid assemblages in the meadow

In congruence with published studies on copepods and nematodes (Novak, 1989; Mascart, Lepoint, et al., 2015), we found that halacarid communities in the leaves consisted mainly of few very abundant species, whereas in the matte, mite abundances were more even between species (Figure 3A-C). We speculate that such difference might be explained by the higher exposure of the leaves to both hydrodynamics (Borg et al., 2006; but see Pugh & King, 1985, in halacarids) and predation (Hovel et al., 2002; Hovel & Fonseca, 2005), which may filter and select for those species that withstand the water currents and avoid predation. In effect, the exposure of the halacarids in the leaves to these stressors might explain that neither their species richness nor abundances were affected by the changes in this habitat (e.g., increase of leaf length and so its epiphytic load; Mabrouk et al., 2010). In contrast, halacarids are protected from predators and water currents in the matte since more individuals occurred in patches of greater habitat complexity. However, unlike other detritivorous groups, such as copepods, annelids, and nematodes (Vizzini et al., 2002; Mirto et al., 2014; Cvitković et al., 2017), halacarids showed no relationship with the organic carbon in the matte. Whether this is due to different dietary preferences in halacarids (Pugh & King, 1986; Bartsch, 1989) will warrant future research.

The findings of our case study supported partially the habitat preferences shown by the literature survey for *P. oceanica* meiofaunal species (Figure 2A and 2D). Indeed, 16 of the 21 halacarid species recorded here co-occurred in both the leaves and the matte. Nevertheless, when accounting for abundances, the species composition of halacarid assemblages differed between the leaves and the matte (Table 2). These differences were mainly due to turnover rather than to differences in species nestedness, indicating a certain habitat sorting between the leaves and matte. Such habitat sorting is evident amongst the dominant species *Rhombognathus praegracilis* and *Copidognathus lamelloides*, preferring the leaves and the matte, respectively (Figure 3E and 3F). Indeed, *Rhombognathus praegracilis* belongs to the phytal-specialist subfamily Rhombognathinae, which possesses morphological adaptations for an epiphytic phytophagous lifestyle, such as complex and thick claws and serrate setae, useful to graze on the rich algal communities of the leaves and withstand the currents (Pugh et al., 1987; Bartsch, 2006; Martínez, García-Gómez, in press). In contrast, *Copidognathus lamelloides,* which thrive in the matte, is attributed with an infaunal lifestyle, as it has been usually reported from sheltered habitats (Somerfield & Jeal, 1995; Barstch, 2009; Riesgo et al., 2010). In addition, juvenile halacarids were significantly rarer than adults in the leaves (Figure 3D). Noticeably, juveniles have developing structures, such as claws, an additional pair of legs, and more leg segments (Bartsch, 2015). These structures enhance the adult’s grip to the substrate, and so, their absence might relegate the juveniles to the matte, which is considerably more protected from currents than the leaves. Overall, our results suggest that, as in nematodes and copepods (Novak, 1989; Mascart, Agusto, et al., 2015), migration between leaves and matte is frequent in halacarids, yet only certain species thrive in each of those habitats.

## Supporting information

Table S1. Records of meiofaunal species in Posidonia oceanica habitats.

Table S2. Summary of the ecological research on meiofauna in Posidonia oceanica.

## Acknowledgements

The authors want to thank Dr Alfonso Ramos-Esplá (CIMAR) for the logistical assistance during the sampling campaigns in Santa Pola. We also thank Dr Sergio Pérez and Nuria Rico for their help with the samplings. We gratefully acknowledge Dr Ilse Bartsch for her invaluable help with the identification of halacarid species. Ilse Bartsch, as well as our colleagues Marco Curini-Galletti, Antonio Todaro, Ulf Jondelius, and Tom Artois, provided unpublished records or helped us find a few records hidden in the old literature. We also thank Dr Dolores Trigo and María Isla for the facilities and their useful assistance in the laboratory of soil chemistry of the Zoology Department (UCM). Thibaud Mascart kindly shared us with the supplementary material of his PhD thesis. Guillermo García-Gómez was supported by an Erasmus+ mobility fellowship, OLS ID 641798. Alejandro Martínez was supported by a European Marie Skłodowska-Curie Actions individual fellowship, project ANCAVE 745530. Nuria Sánchez was funded by the Research Talent Attraction Program for incorporation into research groups in the Community of Madrid (2019-T2/AMB-13328)

## Ethical approval

All applicable international, national, and/or institutional guidelines for the care and use of animals were followed (Directive 2010/63/EU).

## AUTHOR CONTRIBUTION STATEMENTS

GGG, AM surveyed the literature and compiled the datasets for the review. GGG, AGH, NS, AM and FP planned the sampling design of the case study. GGG, AGH, NS and AIM collected the samples; GGG, AGH and NS sorted the latter samples for animals of interest and measured the environmental variables, whereas GGG identified the animals. GGG, AM and DF planned the statistical approach and performed analyses. FP provided facilities and support. GGG, AM and DF wrote the first draft. All authors contributed to the writing to additions and comments to the text.

## SUPPORTING INFORMATION

**Table S1.** Records of meiofaunal species in *Posidonia oceanica* habitats compiled during our review, both collected from the literature as well as directly from the specialists’ unpublished records.

Table S2. Summary of the ecological research on meiofauna in *Posidonia oceanica*.

**Table S3.** Results of the linear models for richness (i.e., number of species), abundance (log_10_-transformed) and evenness (Pielou’s J), and the environmental factors measured in each habitat, reported as type-II analysis-of-variance tables. Abbreviation: Df, Degrees of freedom.

| Habitat | Response variable | Environmental predictors | Df | F | *p* |
| --- | --- | --- | --- | --- | --- |
| Leaves | Richness | length of leaves | 1 | 0.104 | 0.751 |
|  |  | density of leaves | 1 | 0.000 | 0.993 |
|  |  | sampling campaign | 3 | 0.194 | 0.899 |
|  |  | residuals | 18 |  |  |
|  | Abundance | length of leaves | 1 | 0.900 | 0.355 |
|  |  | density of leaves | 1 | 2.481 | 0.133 |
|  |  | sampling campaign | 3 | 9.874 | **0.005** |
|  |  | residuals | 18 |  |  |
|  | Evenness | length of leaves | 1 | 0.006 | 0.941 |
|  |  | density of leaves | 1 | 3.593 | 0.074 |
|  |  | sampling campaign | 3 | 1.487 | 0.252 |
|  |  | residuals | 18 |  |  |
| Matte | Richness | density of matte | 1 | 4.052 | 0.059 |
|  |  | organic matter | 1 | 0.063 | 0.805 |
|  |  | sampling campaign | 3 | 3.103 | 0.053 |
|  |  | residuals | 18 |  |  |
|  | Abundance | density of matte | 1 | 9.232 | **0.007** |
|  |  | organic matter | 1 | 0.254 | 0.620 |
|  |  | sampling campaign | 3 | 3.153 | 0.050 |
|  |  | residuals | 18 |  |  |
|  | Evenness | length of leaves | 1 | 0.207 | 0.655 |
|  |  | density of leaves | 1 | 0.810 | 0.381 |
|  |  | sampling campaign | 3 | 0.006 | 0.999 |
|  |  | residuals | 17 |  |  |

